# CYRI-B loss promotes enlarged mature focal adhesions and restricts microtubule and ERC1 access to the cell leading edge

**DOI:** 10.1101/2024.03.26.586838

**Authors:** Jamie A. Whitelaw, Sergio Lilla, Savvas Nikolaou, Luke Tweedy, Loic Fort, Nikki R. Paul, Sara Zanivan, Nikolaj Gadegaard, Robert H. Insall, Laura M. Machesky

## Abstract

CYRI proteins promote lamellipodial dynamics by opposing Rac1-mediated activation of the Scar/WAVE complex. This activity also supports resolution of macropinocytic cups, promoting internalisation of surface proteins, including integrins. Here, we show that CYRI-B also promotes focal adhesion maturation and dynamics. Focal adhesions in CYRI-B-depleted cells show accelerated maturation and become excessively large. We probed the composition of these enlarged focal adhesions, using a Bio-ID screen, with paxillin as bait. Our screen revealed changes in the adhesome suggesting early activation of stress fibre contraction and depletion of the integrin internalisation mediator ERC1. Lack of CYRI-B leads to more stable lamellipodia and accumulation of polymerised actin in stress fibres. This actin acts as a barrier to microtubule targeting for adhesion turnover. Thus, our studies reveal an important connection between lamellipodia dynamics controlled by CYRI-B and microtubule targeting of ERC1 to modulate adhesion maturation and turnover.

## Introduction

As cells migrate over planar surfaces, they create broad, flat membrane protrusions at the front, termed lamellipodia. Activation of the small GTPase Rac1 triggers actin assembly in lamellipodia through binding to the Scar/WAVE complex subunit CYFIP1 (Chen et al., 2010). Binding to Rac1 allows conformational changes of the complex and activation of the Arp2/3 complex to nucleate a branched actin filament network providing the protrusive forces required to extend the plasma membrane (Mullins et al., 1998). The cell’s connection to the surrounding extracellular matrix (ECM) guides migration of individual cells and in multi-cellular organisms, underpinning fundamental processes such as embryogenesis and cancer metastasis. There have been many different types of cell-ECM adhesions described, such as focal complexes, focal adhesions, fibrillar adhesions and 3D matrix adhesions (Doyle et al., 2022). However, they all share a common characteristic that the engaged integrins connect to the actin cytoskeleton through a complex of core adhesion proteins (Geiger et al., 2009). Engaged integrins allow the cell to sense and respond to the surrounding environment by converting mechanical stimuli from focal adhesions to biochemical signals, in a process commonly known as mechanotransduction (Humphrey et al., 2014).

Focal adhesions (FAs) form by the engagement of integrins to the matrix along the cell periphery at the lamellipodia tip (Giannone et al., 2007; Zaidel-Bar et al., 2003). Initially adhesions resemble small dot-like structures known as nascent adhesions, which mature and enlarge, changing in protein composition. Over 2000 proteins have been identified as enriched in fibronectin-induced adhesions, but a core of 60 proteins that have been most commonly identified is known as the core adhesome (Horton et al., 2016). Paxillin is one of the earliest proteins recruited to nascent adhesions and is associated with signalling pathways such as via focal adhesion kinase (FAK) through its two binding sites at the N-terminal domain (Legerstee and Houtsmuller, 2021; Scheswohl et al., 2008). FAK is responsible for the recruitment of talin to the nascent adhesions which links the cytoplasmic tails of integrins to the actin cytoskeleton (Lawson et al., 2012). This in turn can influence FA size, which links to cell migration speeds (Kim and Wirtz, 2013) and is reported as a measure of integrin signalling during epithelial-mesenchymal transition (EMT) in many cell types (Legerstee and Houtsmuller, 2021; Tsubouchi et al., 2002). Phosphorylation of integrin-mediated adhesions by paxillin and FAK activates the small GTPase Rac1 in a signalling cascade, which in turn activates the Scar/WAVE complex and enhances membrane protrusion (Zaidel-Bar et al., 2005). As the cell moves forward, the nascent adhesions become associated with the lamellipodium-lamellum interface (Alexandrova et al., 2008), where the retrograde flow rate reduces, and adhesions either disappear or enlarge into mature focal adhesions engaged with actin bundles. These recruit additional adaptor and signalling proteins such as vinculin, zyxin and α-actinin and begin to exert mechanical forces upon the actin cytoskeleton (Burridge and Guilluy, 2016; deMali et al., 2002). Maturation is a positive feedback loop, triggering further clustering of activated integrins (Humphries et al., 2007) to strengthen the actin-integrin connections, elongation and strengthening of, links with contractile actin stress fibers containing myosin-II (Pellegrin and Mellor, 2007).

As cells migrate, FAs linked to the ECM disassemble and the disengaged integrins are internalised and degraded or recycled back to the plasma membrane (Moreno-Layseca et al., 2019). This can be facilitated by the protease calpain cleaving integrins and talin (Franco et al., 2004; Kerstein et al., 2017), dynamin and clathrin-mediated endocytosis and clathrin-independent mechanisms such as macropinocytosis and caveolin-mediated endocytosis (Maritzen et al., 2015). Membrane trafficking and microtubules play an important dual role in FAs, both in positive trafficking of integrins to nascent adhesions and in trafficking of relaxation or disassembly factors such as metalloproteases to degrade matrix (Garcin and Straube, 2019; Seetharaman and Etienne-Manneville, 2019; Stehbens et al., 2014). Microtubules are also thought to promote endocytosis at focal adhesions, possibly mediating integrin internalisation (Ezratty et al., 2005). To enhance FA turnover, microtubules are targeted to FA sites by CLASP-mediated capture to the ends of actin stress fibers via a complex of proteins including LL5β, ERC1 and Liprin-α1 (Astro et al., 2014; Astro et al., 2016; Lansbergen et al., 2006; Stehbens et al., 2014). These in turn link to talins via the adaptor Kank proteins to release the FA complex of proteins on the cytoplasmic side (Bouchet et al., 2016; Paradzik et al., 2020). ERC1 targeting promotes the internalisation and recycling of surface integrins (LaFlamme et al., 2018) via Rab7-dependent vesicles along microtubules (Astro et al., 2016).

Lamellipodia and adhesion dynamics are fundamental for cell behaviour. We recently showed that loss of the Scar/WAVE complex by *NckAP*1 deletion had a negative effect on FA turnover and cell migration (Whitelaw et al., 2020). Furthermore, the Scar/WAVE complex has been implicated in the internalisation and recycling of integrins (Rainero et al., 2015). Recently, we identified a novel class of Rac1 interacting proteins that act as negative regulators of the Scar/WAVE complex activation, termed CYFIP-related RAC1 interacting (CYRI) proteins (Fort et al., 2018). There are two isoforms of CYRI proteins in mammals, named CYRI-A and CYRI-B for the genes (*CYRIA*, *CYRIB (human) and Cyri-a, Cyri-b (mouse)*, formerly known as *FAM49A, Fam49a* and *FAM49B, Fam49b*, respectively. CYRI proteins oppose Rac1-mediated activation of Scar/WAVE and Arp2/3 and thus control cell migration and chemotaxis (Fort et al., 2018), macropinocytic structures (Le et al., 2021) and pathogen invasion (Yuki et al., 2019). Here we show that deletion of *Cyri-b* enhances FA assembly during early stages of spreading and alters the recruitment of core FA proteins. FAs become larger and more mature in *Cyri-b* KO cells than controls. We performed a Bio-ID screen to detect changes in composition of FAs in CYRI-B depleted cells. Among the changes, we found that *Cyri-b* KO cells have reduced ERC1 in the vicinity of paxillin by proximity biotinylation and at the leading edge, by immunofluorescence. This paucity of ERC1 is accompanied by reduced microtubule recruitment to the cell periphery, likely promoting the stable enlarged FAs by preventing microtubule-stimulated turnover.

## Results

### Focal adhesions are elongated and larger in *Cyri-b* KO cells

CYRI-B restricts lamellipodia spreading and directed cell migration by dynamically sequestering active Rac1 away from the Scar/WAVE complex (Fort et al., 2018). Nascent adhesions form within the lamellipodia region of migrating cells and coupled with the actin retrograde flow, mature into FAs (Hu et al., 2007). Therefore, we asked how loss of CYRI-B might affect FAs. We deleted *Cyri-b* in B16-F1 mouse melanoma cells using transient CRISPR-Cas9-GFP (Ran et al., 2013). Cas9-GFP positive B16-F1 cells were sorted by flow cytometry and the clones were tested for the loss of CYRI-B by Western blotting (Fig. S1a). As previously reported (Fort et al., 2018), *Cyri-b* knockout (KO) clones in B16-F1 cells spread rapidly (Fig. S1b) and formed large, broad lamellipodia (Fig. 1a,b). For this study, we focused on clone #3 and confirmed the deletion of *Cyri-b* by immunoblot (Fig. 1a, S1a).

**Figure 1:**
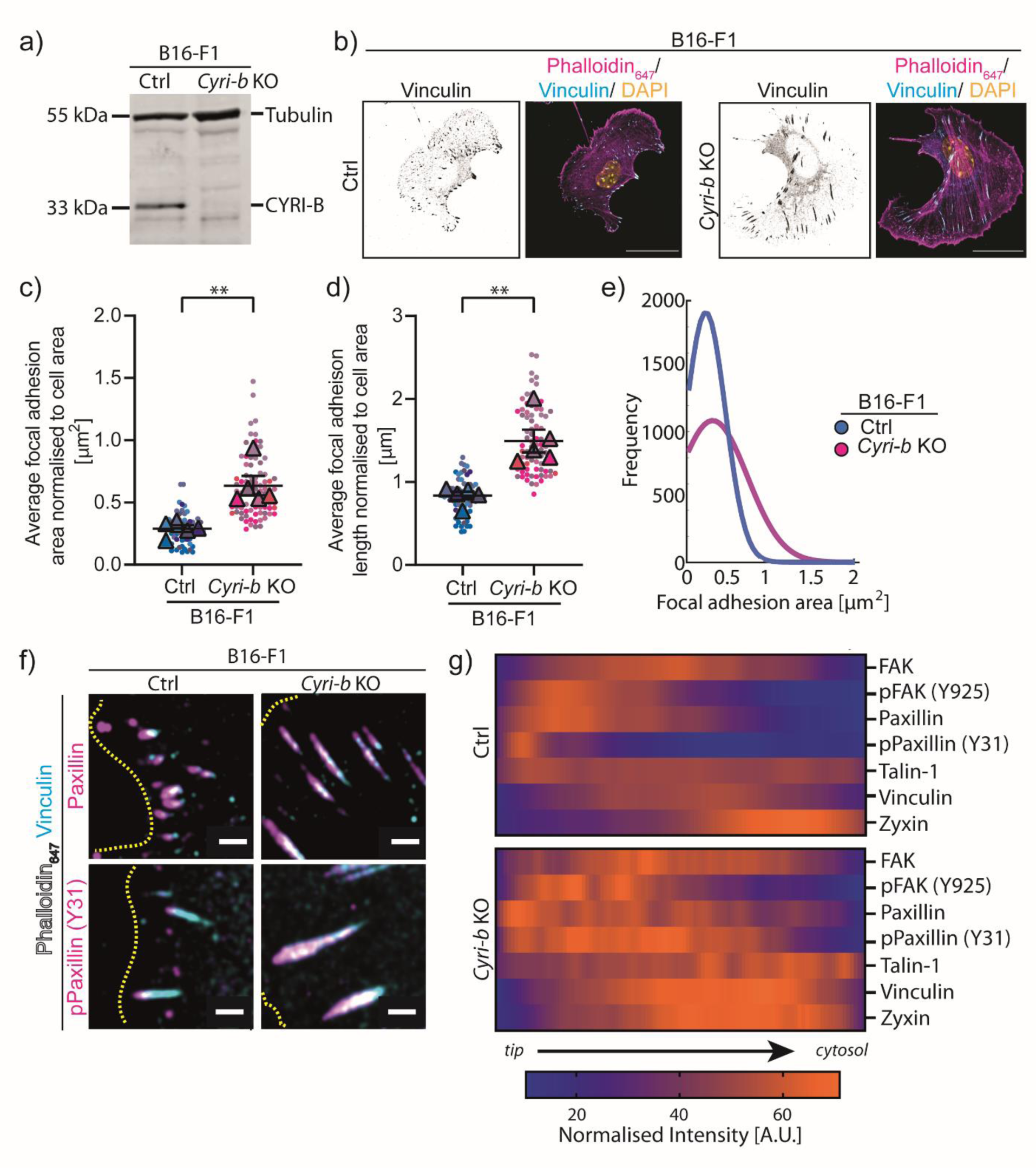
Focal adhesions are elongated and show enhanced phospho-paxillin in *Cyri-b* knockout cells. **a)** Immunoblot of CRISPR-Cas-9 knockout of *Cyri-b* in B16-F1 cells. Tubulin as loading control and anti-CYRI-B. **b)** FA sizes were compared in B16-F1 Ctrl and *Cyri-b* KO cells. Representative images B16-F1 cells spreading on laminin-coated coverslips and stained with vinculin (Cyan), phalloidin (Magenta) and DAPI (Yellow). Greyscale image of vinculin on the left. Scale bar 25 µm. FA area **c)** or FA length **d)**. A total of 69 control and 79 *Cyri-b* KO cells were analysed from 5 independent experiments. Superplots analysed with n=5 and a paired parametric t -test. ** P-value <0.01. **e)** An independent analysis of FA area detected by CellProfiler and presented as a line distribution of the frequency. **f-g)** Comparisons of FA composition between B16-F1 control and *Cyri-b* KO cells using vinculin antibodies to normalise. **f)** Representative images with vinculin (cyan) and the comparative FA antibody (magenta). The leading edge of the cell is highlighted by a dashed yellow line. Scale bars represent 2 µm. **g)** Profiles of FAs were measured with the intensity normalised to the corresponding vinculin intensity. The colour heat map indicates the average intensity of FA proteins from the FA tip through to the end facing the cytoplasm. Orange represents a high fluorescence intensity e.g. strong localisation. Purple represents low fluorescence intensity indicating weak localisation within the FA.

Loss of CYRI-B resulted in large, elongated focal adhesions spread throughout the lamellipodium and cell body of B16-F1 cells (Fig. 1b-e). Quantification of FA area using CellProfiler showed that the *Cyri-b* KO cells had an increased frequency of larger FAs (Fig. 1e). We confirmed the enlargement of FAs in *Cyri-b^fl/fl^*mouse embryonic fibroblasts (MEFs) with Cre-ERT2 (Fort et al., 2018), which deletes *Cyri-b* upon addition of 4-hydroxytamoxifen. MEFs generally displayed larger FAs than B16 F1 cells, but these were further enlarged upon deletion of *Cyri-b* (Fig. S1c-d).

To explore maturation status of the larger FAs in CYRI-B depleted cells, we probed the distribution of key protein components of the adhesion machinery. By creating a heat map of the intensities of each protein and averaging this over several FAs (Fig. 1f), measuring from the most peripheral point (tip) towards the cell centre (cytosol) (Fig. S1d), we compared the distributions of FAK, paxillin, talin-1 and zyxin to that of vinculin (Fig. 1f,g, Fig. S1e-g). Paxillin displays a similar profile in the control (Ctrl) and *Cyri-b* KO cells but shows a broader distribution in the *Cyri-b* KO cells. There was also a large increase in the intensity and breadth of phospho-paxillin (Y31), which has been shown to be important for cell migration (Petit et al., 2000) (Fig. 1g, Fig. S1f). The distribution of FAK was similar between Ctrl and *Cyri-b* KO cells (Fig. 1g, Fig. S1f). We also checked the phosphorylation of FAK Tyr-925 due to its role in cell migration through its activation of the p130Cas/Rac1 signalling pathways (Deramaudt et al., 2011). However, similar to FAK, pFAK^Y925^ showed only slight changes in distribution (Fig. 1g, Fig. S1f).

Talins directly connect to both integrins and F-actin (Das et al., 2014; Jin et al., 2015), while vinculin is recruited to talin and reinforces the F-actin anchoring (Bays and DeMali, 2017; Boujemaa-Paterski et al., 2020). As expected, vinculin and talin localisation span the whole FA in both the control and *Cyri-b* KO cells (Fig. S1f). However, of note, talin-1 exhibits prominent intensity peaks to the rear half of the FA in the *Cyri-b* KO cells that are not observed in the control cells (Fig. 1g, Fig. S1f). Furthermore, while vinculin is spread throughout the FAs similarly in Ctrl and *Cyri-b* KO cells, the intensity of vinculin is greater in the *Cyri-b* KO cells and zyxin is similar but has a broader distribution in the *Cyri-b* KO cells. (Fig. 1g, Fig. S1f). In summary, FAs in CYRI-B depleted cells show enhanced phospho-paxillin and enhanced recruitment of several other core FA proteins, suggesting that the larger FAs are more mature, which might reflect reduced turnover dynamics.

We next examined how the larger FAs in *Cyri-b* KO cells formed and matured over time. B16-F1 cells were replated and fixed at different time points during adhesion to observe a time progression from early focal complex formation to more mature FAs (Geiger et al., 2009). We used paxillin as a marker of early focal complex formation, which we expected to remain through to FA maturity and also zyxin as a marker for mature FAs (Legerstee and Houtsmuller, 2021). *Cyri-b* KO cells recruited proteins such as paxillin to the focal complex as early as 30 minutes and the adhesion sizes quickly increased within the 3-hour time-course (Fig. 2a-c). Similarly, zyxin was also observed in the FAs after 30 minutes (Fig. 2a-c), indicating that even these early focal complexes displayed markers of mature FAs (Fig. 2b-c). Control cells took around 30 minutes longer to form discernible FAs (Fig. 2a-c).

**Figure 2:**
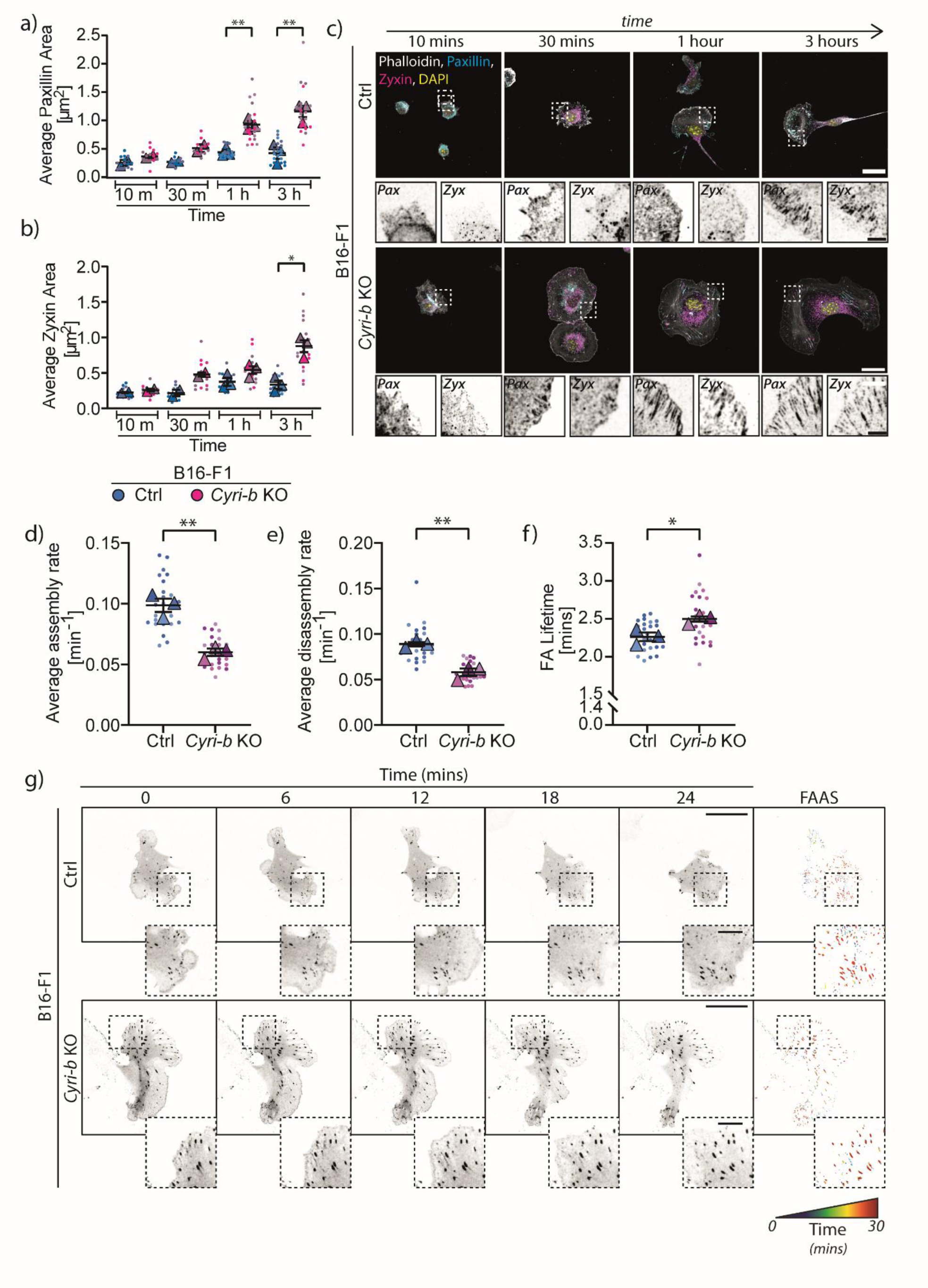
Adhesion dynamics are altered in the *Cyri-b* KO cells. **a-c)** The formation and maturation of FAs in B16-F1 cells from initial seeding to cells spreading. Phalloidin (white) was used as a marker for the cell size, Paxillin (Cyan) was used as an early FA marker and Zyxin (Magenta) was used as a later marker for mature FAs. Cells were trypsinised and seeded for the indicated time before fixation. **a)** The average paxillin area and **b)** the average zyxin area over time for the control and *Cyri-b* KO cells. 15 cells from 10-30 minutes and 25 cells for 1-3 hours analysed from ≥2 independent experiments. Mean ± S.E.M., two-tailed paired t-test comparing control and *Cyri-b* KO cells on n=2 (10 and 30 minutes) or n=3 (1 and 3 hours) experiments in Superplot format. * P<0.05, ** P<0.01. **c**) Representative images for the time course experiment. Scale bar represents 25 μm and the inset 2.5 μm. **d-g**) focal adhesion dynamics of 27 cells from 3 independent experiments. Cells expressing pEGFP-Paxillin were assessed for their focal adhesion assembly rates (**d**) and disassembly rates (**e**). **f**) The lifetimes of the focal adhesions. Error bars represent Mean ± S.E.M. in superplot format. Statistical differences determined by a two-tailed paired t-test comparing control and *Cyri-b* KO cells, * P<0.05, ** P<0.01. **g**) Representative images of focal adhesion turnover over the 30-minute time course. For the FAAS, there adhesions are colour coded through time from blue at the start to red at the end of the experiment. Scale bar represents 25 μm and 5 μm for inset.

This was followed by investigating the dynamics of the large FAs in the *Cyri-b* KO compared to the control B16-F1 cells by measuring the assembly and disassembly rates and the lifetime of the adhesions after the cells had been allowed to attach and migrate in a steady state. Live imaging of the cells expressing pEGFP-Paxillin were captured over a 30-minute time course and analysed using the Focal Adhesion Analysis Server (FAAS) (Berginski and Gomez, 2013). Here we observed that the adhesions in the control cells were able to form and disassemble much faster than those in the *Cyri-b* KO cells (Fig. 2d,e,g; Supp. Movie1). It was apparent when calculating the longevity of the adhesions that those in the *Cyri-b* KO persisted for longer (Fig. 2f). Overall, this indicates that these large FAs in the *Cyri-b* KO are more stable than those of the control cells.

### The large focal adhesions in *Cyri-b* KO cells are not solely due to increased Rac1 activity

During spreading, α5β1 integrin signalling leads to Rac1 activation at the leading cell edge and subsequent lamellipodia protrusions (Price et al., 1998). This increases FAK and paxillin phosphorylation leading to increased activation of the p130Cas/Dock180/Rac1 pathways in a positive feedback loop (Valles et al., 2004). As growth of the adhesions progresses, Rac1 is replaced by RhoA, activating contractile forces along the FAs (Arthur and Burridge, 2001). Loss of CYRI-B causes the cells to form large lamellipodia due to an increased activity of active Rac1, inducing Scar/WAVE activity (Fort et al., 2018). We speculated that increased Rac1 activity in the *Cyri-b* KO could be enhancing the formation and maturation of FAs. To test this, we expressed constitutively active mutant Rac1^Q61L^-GFP into B16-F1 wild-type (WT) cells. FA sizes were significantly larger in Rac1^Q61L^- GFP expressing cells, but importantly these FAs were still significantly smaller than those of *Cyri-b* KO cells (Fig. 3a-c). We also rescued the *Cyri-b* KO cells with CYRI-B-p17-GFP an internally tagged CYRI construct in which GFP is inserted after residue 17 of CYRI-B (Le et al., 2020). CYRI-B-p17-GFP rescue restored normal FA sizes. We rescued with CYRI-B^R160/161D^-p17-GFP, a construct with mutations preventing Rac1 interaction (Fort et al., 2018), which conferred a reduction of FA sizes but only to a level similar to cells expressing Rac1^Q61L^ (Fig.3a-c). Overall, this suggests that increased Rac1 activity in the *Cyri-b* KO cells only partially contributes to the large FA size.

**Figure 3:**
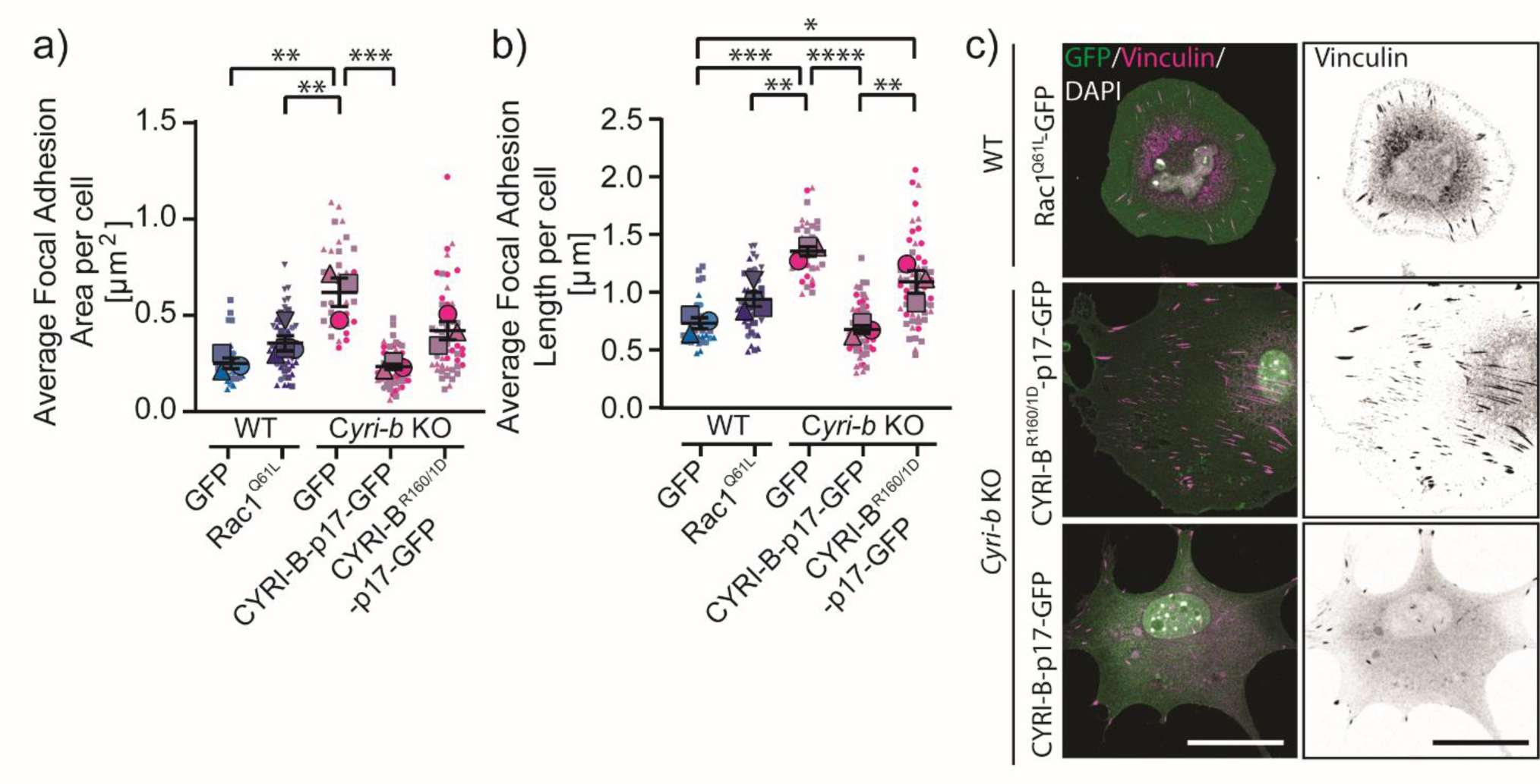
Increased Rac1 activity alone does not account for the enlarged focal adhesions in *Cyri-b* knockout cells. **a-c)** FA sizes in B16-F1 cells expressing different GFP constructs to assess whether increased Rac1 is activity is responsible for the large FAs in the *Cyri-b* KO cells. B16-F1 WT cells expressing pEGFP-Rac1^Q61L^ or *Cyri-b* KO cells rescued with CYRI-B-p17-GFP or CYRI-B^R160/1D^-p17-GFP (Rac1 binding mutant). **a)** FA area. **b)** FA length. 35 WT + GFP only, 53 WT + Rac1^Q61L^-GFP, 35 *Cyri-b* KO, 56 *Cyri-b* KO + CYRI-B-p17-GFP and 57 *Cyri-b* KO + CYRI-BR160/1D-p17-GFP cells analysed from 3 independent experiments, shown by the different symbols. Error bars represent mean ± S.E.M., 1-way ANOVA on n=3 independent experiments in superplot format. * P<0.05, ** P<0.01, *** P<0.001, **** P<0.0001. **c)** Representative images of FA in cells expressing GFP fusion constructs. Left hand side shows merge with GFP (green), vinculin (magenta) and DAPI (white). Right side shows images of vinculin in greyscale. Scale bar 25 μm.

### BioID screen for Paxillin interactions reveals altered focal adhesion networks in *Cyri-b* KO cells

To identify additional factors that might affect FA maturation dependent on CYRI-B, we used paxillin as the bait in a proximity biotinylation Bio-ID experiment (Dong et al., 2016) (Fig. S2a,b). Proximity biotinylation of paxillin was previously used to provide insight into the molecular composition of FAs to define the adhesome (Chastney et al., 2020; Dong et al., 2016). Indeed, our Bio-ID screen identified enrichment of well-known FA proteins such as talin-1, -2, FAK, adhesion regulators such as Kank2, small GTPase interactors such as GIT1 and β-PIX and actin-binding proteins such as Shroom2, 4 in the larger FA of *Cyri-b* KO cells (Fig. 4a, Fig. S2c,d,f). Interestingly, zyxin, a protein found in more mature FA, was enriched in the *Cyri-b* KO adhesions compared to the control cells, reconfirming the idea that the FAs in the *Cyri-b* KO cells are more mature and in agreement with our immunofluorescence analysis (Fig. 1g, 2b). On the other hand, the cytoskeleton and membrane trafficking adaptor protein, ERC1 was depleted in the proximity of adhesions of *Cyri-b* KO cells (Fig. 4a). ERC1 mediates displacement of cytoplasmic adhesion complex proteins, thus promoting the internalisation of surface integrins via clathrin-mediated and clathrin-independent endocytosis (Astro et al., 2016; Pellinen et al., 2006).

**Figure 4:**
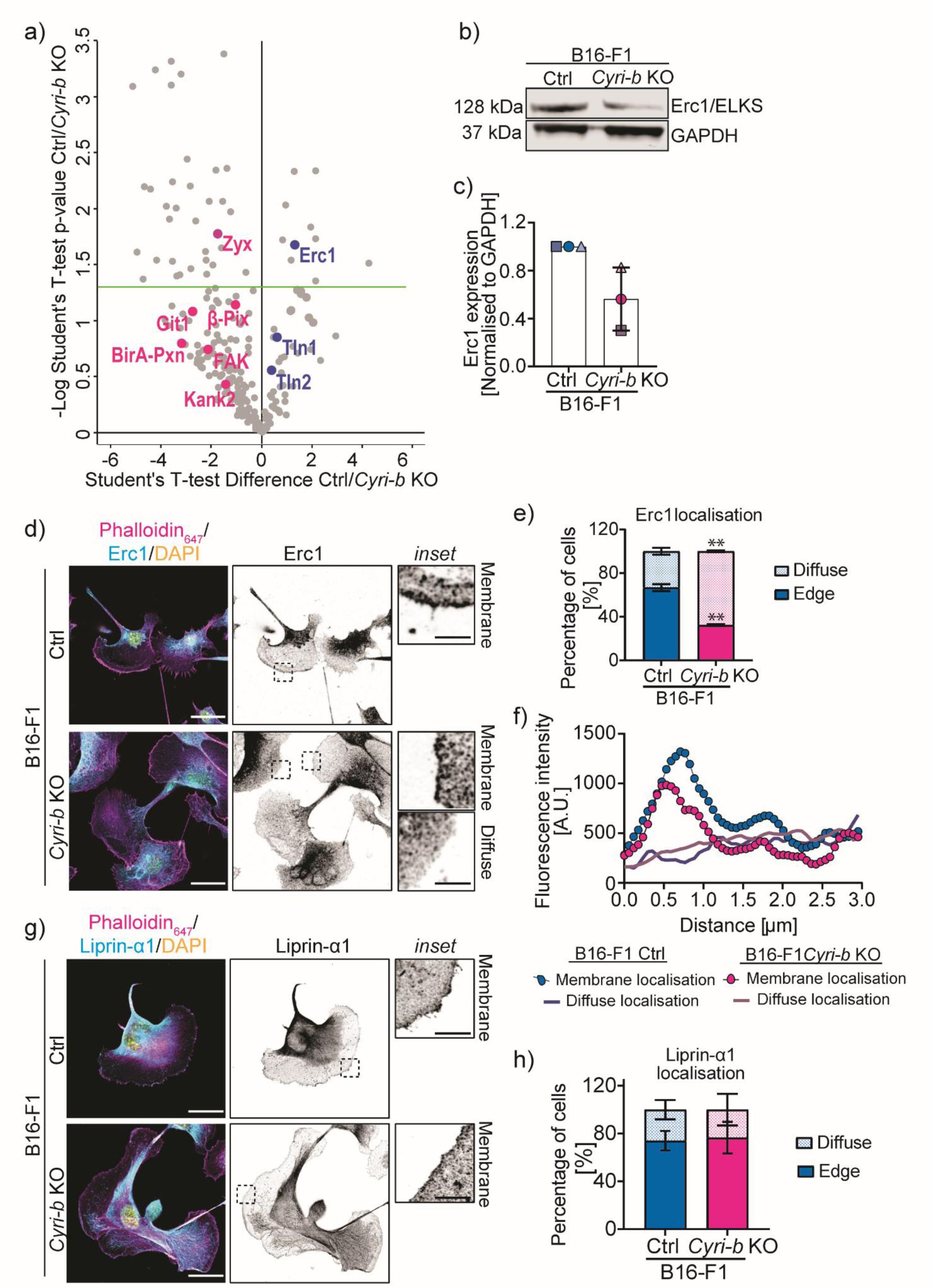
BioID screen of Paxillin reveals decreased association with ERC1 in *Cyri-b* knockout cells. **a)** Volcano plot displaying the results from the proximity biotinylation screen of paxillin in B16 -F1 control and *Cyri-b* KO cells. Proteins enriched in proximity to paxillin in *Cyri-b* KO cells are shown in magenta and proteins enriched in control cells shown in blue. Proteins above the green horizontal line are enriched in either control or *Cyri-b* KO cells. See also Figure S2 for details of other enriched proteins. P-value <0.05. **b)** Representative Western blot of endogenous ERC1 levels in B16-F1 control and *Cyri-b* KO cells. **c)** Quantification of ERC1 from western blotting normalised to GAPDH loading control. Error bars represent Mean ± S.D. from three independent experiments. **d)** Representative images of ERC1 localisation using an anti-ERC1 antibody. Actin cytoskeleton (magenta), ERC1 (cyan) and DAPI (yellow). Insets depict ERC1 localisation either at the membrane or as a diffuse cytosolic staining. Scale bars represents 25 μm and 5 μm for inset. **e)** Quantification of ERC1 localisation to cell edge (solid colour) or diffuse in the cytoplasm (coloured dots). 61 control and 65 *Cyri-b* KO cells analysed from 3 independent experiments and converted to percentages. Mean ± S.D., two-tailed t-test with Welch’s correction. ** P<0.01 **f)** Fiji plot profile fluorescence intensity of the localisation of ERC1 staining. The lines with circles represent the average intensity of the ERC1 signal from cells with a membrane localisation. The lines without circle points represents the intensity of diffuse staining showing a lack of intensity at the membrane. The distance measured was 3 μm from the leading edge into the cell. n=19 control and 19 *Cyri-b* KO cells. **g)** Representative images of Liprin-α1 localisation using an anti-Liprin-α1 antibody (cyan), actin cytoskeleton (magenta) and DAPI (yellow). Liprin-α1 channel displayed in greyscale to the right-hand side. Scale bars represents 25 μm and 5 μm for inset. Insets depict Liprin-α1 localisation at the membrane. **h)** Quantification of Liprin-α1 at the plasma membrane (solid colour) vs diffuse cytoplasmic staining (coloured dots). 42 control and 54 *Cyri-b* KO cell analysed from 3 independent experiments and converted to percentages. Mean ± S.D. Two-tailed t-test with Welch’s correction with no significance reached.

### ERC1 but not Liprin-α1 is affected by the loss of CYRI-B

Due to its importance in integrin internalisation, we investigated ERC1 depletion at *Cyri-b* KO FA further. Immunoblotting showed that that ERC1 total protein levels are reduced in *Cyri-b* KO B16-F1 cells (Fig. 4b,c). Moreover, ERC1 is thought to form a complex with Liprin-α1 and LL5β and localise to the leading edge of migrating cells (Astro et al., 2014). ERC1 has a clear localisation to the leading edge in around 70 % of control cells but this was reduced to around 30 % in *Cyri-b* KO cells (Fig. 4d,e). Moreover, localisation of ERC1 at the leading edge of *Cyri-b* KO cells was tighter, with a reduced fluorescence intensity (Fig. 4f). Conversely, Liprin-α1 (LAR-interacting protein 1), the complex partner of ERC1 which marks synaptic vesicle docking sites in neuronal cells (Astro et al., 2016; Ko et al., 2003; Liang et al., 2021), localised to the leading edge in approximately 70% of both the control and *Cyri-b* KO cells (Fig. 4g,h), suggesting that ERC1 depletion is relatively specific following the loss of CYRI-B and in line with a previous study showing that Liprin-α1 localisation does not depend on ERC1 (Astro et al., 2016). To ask whether ERC1 interacted with CYRI-B directly, we performed a GFP-trap experiment with GFP-CYRI-B and probed for endogenous ERC1, however we did not detect any interaction (Fig. S2e). This suggests that the effect of CYRI-B depletion on ERC1 localisation is likely to be indirect.

We reasoned that if loss of *Cyri-b* affects FA size via a reduced association of ERC1 with FAs, then depletion of ERC1 should enhance FA size. Using a pool of small interfering RNAs (siRNA) specific to *Erc1*, we achieved a greater than 70% reduction in ERC1 protein levels (Fig. 5a,b). B16-F1 cells depleted of ERC1 resembled *Cyri-b* KO cells (Fig. 1c), displaying larger cell area (Fig. 5c) and large elongated FAs (Fig. 5d-f). This supports our hypothesis that loss of *Cyri-b* affects adhesions and spreading at least partly via interfering with ERC1 recruitment to FAs, which in turn affects FA dynamic turnover.

**Figure 5:**
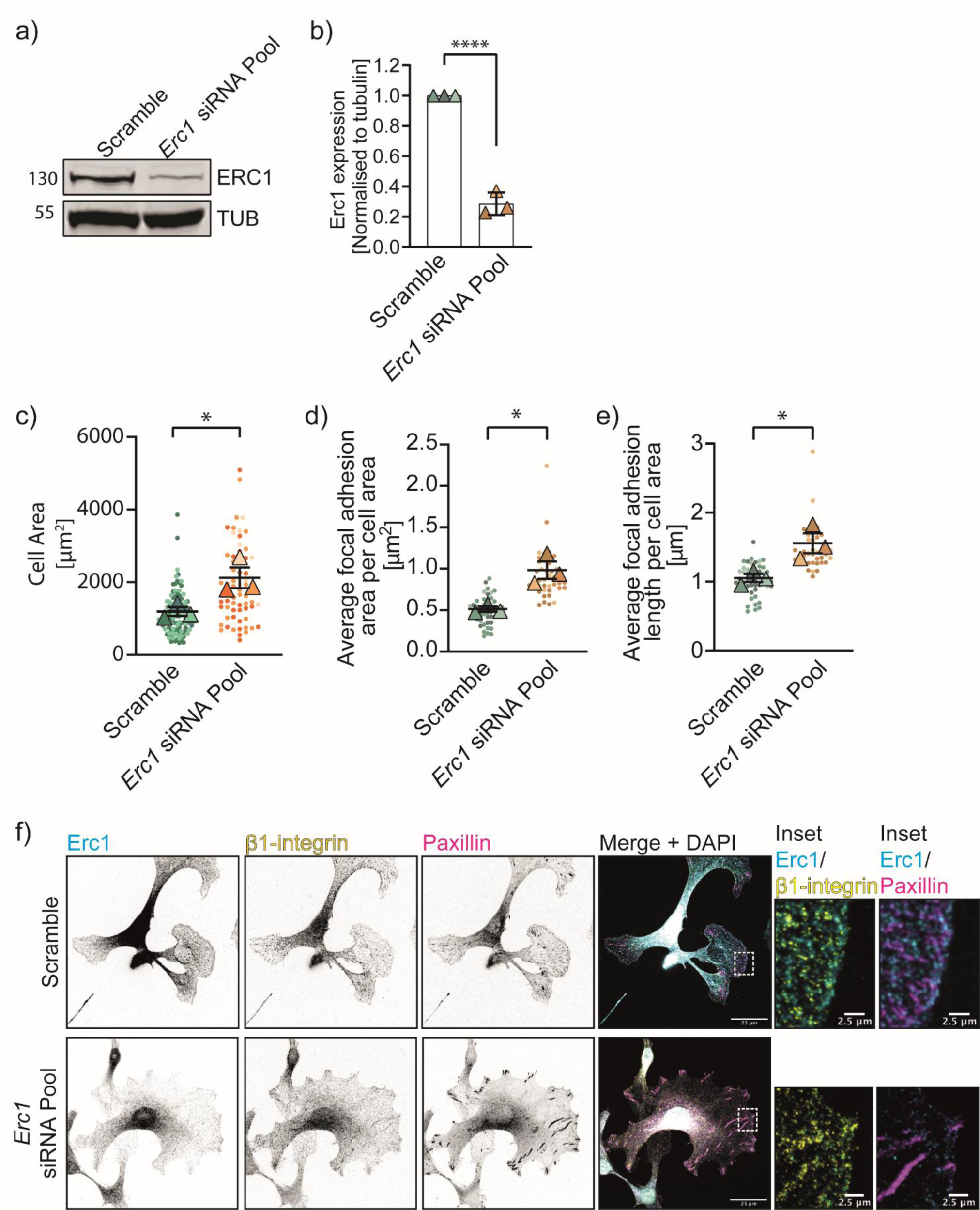
Depletion of ERC1 increases focal adhesion sizes. Downregulation of *Erc1* in B16-F1 cells using 10 nM specific siRNAs pooled. **a)** Representative western blot of ERC1 levels in either B16-F1 cells treated with a scramble or pooled siRNA against Erc1.Tubulin (TUB) as loading control. **b)** Western blot quantification of ERC1 levels in B16-F1 scramble or ERC1 siRNA pool from 3 independent experiments. Mean ± S.D. Two-tailed t-test. **** P<0.0001. **c)** Cell area of B16-F1 scramble or *Erc1* siRNA pool. 106 scramble and 63 cells analysed from 3 independent experiments. Mean ± S.E.M., two-tailed paired t-test on the independent averages from n=3 experiments in superplot format. * P<0.05. **d)** Average FA area per cell **e)** Average FA length. **d-e)** 42 scramble and 42 ERC1 knockdown cells analysed from 3 independent experiments. Mean ± S.E.M., two-tailed paired t-test on the independent average from n=3 experiments in superplot format. * P<0.05. **f**) Representative images ERC1 (cyan), β1-integrin (yellow) or paxillin (magenta). Scale bar 25 μm for main image and 2.5 μm for inset.

### Loss of *Cyri-b* or ERC1 similarly impairs integrin internalisation

Depletion of ERC1 was previously linked to a reduction of internalised β1-integrin receptors and reduced lamellipodial persistence and migration (Astro et al., 2014). We hypothesised that the reduced ERC1 expression in the *Cyri-b* KO cells may increase β1-integrin display at the cell surface. Indeed, we detected an increase in β1-integrin focal adhesion area on the surface of migrating *Cyri-b* KO B16-F1 cells (Fig. 6a,b) that was comparable to what we observed for other FA markers (Fig. 1c). We also observed a 2-fold increase in total β1-integrin levels in *Cyri-b* KO cells (Fig. 6c).

**Figure 6:**
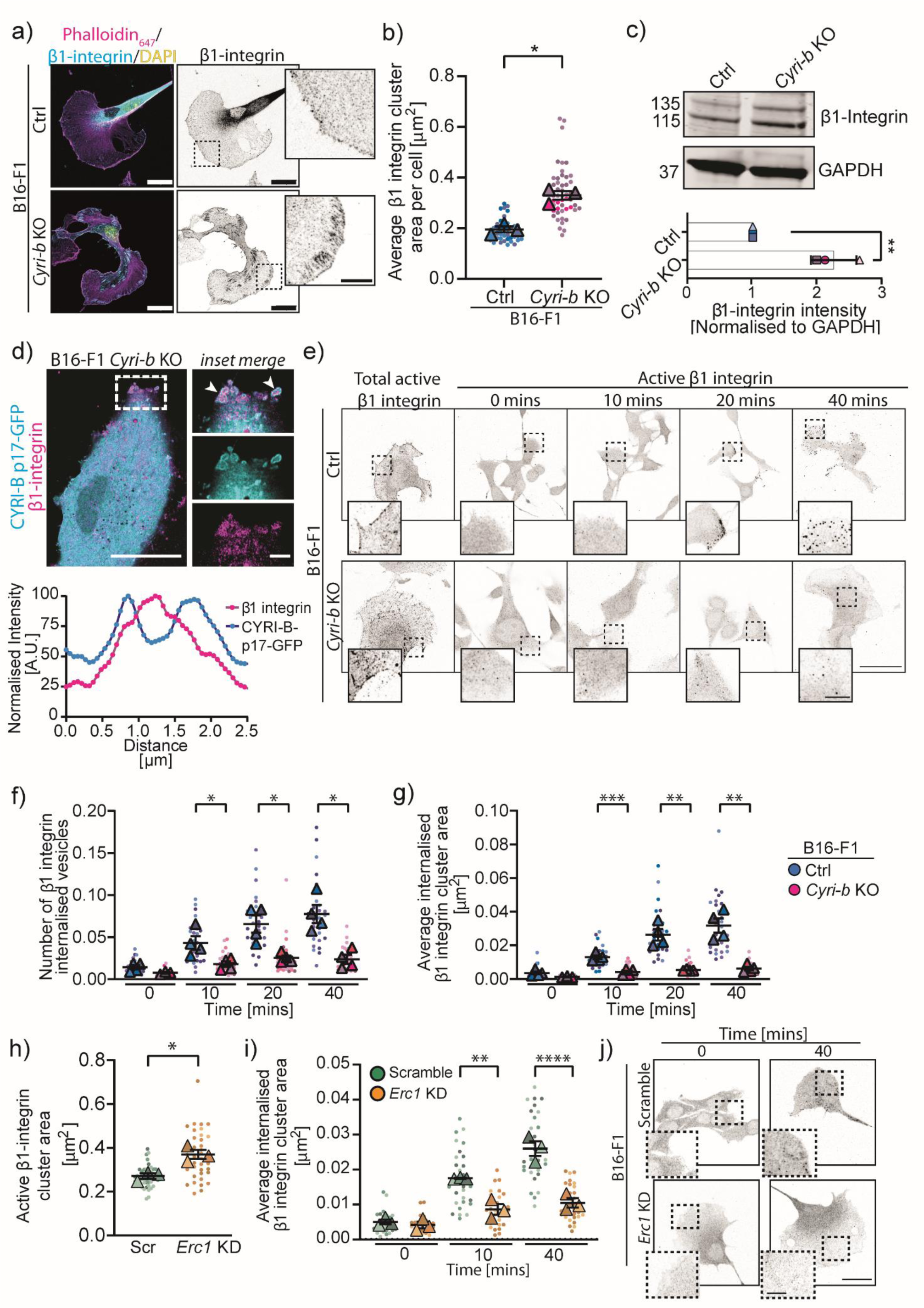
Loss of *Cyri-b* or *ERC1* reduces integrin internalisation. **a)** Immunofluorescence images of β1-integrin staining (cyan), actin cytoskeleton (magenta) and DAPI (yellow). Right-hand image; β1-integrin staining in greyscale. Scale bars represents 20 μm and 5 μm for insets **b)**. Quantification of the average β1-integrin cluster area in B16-F1 control and *Cyri-b* KO cells. 45 control and 50 *Cyri-b* KO cells analysed from 3 independent experiments. Mean ± S.E.M., two-tailed paired t-test on n=3 experiments in superplot format. * P<0.05. **c)** Western blot and quantification of β1-integrin levels in control and *Cyri-b* KO cells from 3 independent experiments. GAPDH as loading control. Unpaired t-test, ** P<0.01. **d)** Live imaging of B16-F1 *Cyri-b* KO cells rescued with CYRI-B-p17-GFP (cyan) and β1-integrin-mCherry (magenta). *Inset,* white arrowheads highlight β1-integrin positive structures surrounded by CYRI-B. Scale bars represents 25 μm and 5 μm for inset. Plot profile of these β1-integrin containing structures shows two peaks of CYRI-B signal intensity (cyan) around the peak of β1-integrin intensity (magenta). **e-g)** β1-integrin internalisation comparison between B16-F1 control and *Cyri-b* KO cells. **e)** Representative images of internalised β1-integrin. Total active β1-integrin characterises the normal β1-integrin localisation within the cells prior to the assay. Time course of β1-integrin internalisation before an acid wash to remove any extracellular bound antibody. Scale bars represents 25 μm and and 5 μm for inset. **f)** Number of β1-integrin internalised vesicles, **g)** average internalised β1-integrin cluster area normalised to cell area over time normalised to cell area. **f-g)** n=30 cells for each condition analysed from 4 independent experiments. 1-way ANOVA on n=4 independent experiments in superplot format. * P<0.05, ** P<0.01, *** P<0.001. **h)** Active β1-integrin cluster area between the scramble control and *Erc1* siRNA KD. 42 scramble and 42 *Erc1* knockdown cells analysed from 3 independent experiments **i)** Average β1-integrin internalised between scramble control and *Erc1* siRNA KD. 30 scramble and 30 *Erc1* knockdown cells analysed from 3 independent experiments. **h-i)** Mean ± S.E.M., two-tailed paired t-test on the independent average from n=3 experiments in superplot format. * P<0.05, ** P<0.01, **** P<0.0001. **j)** Representative images of internalised β1-integrin internalisation in B16-F1 scramble or *Erc1* KD cells. Time scale at 0 and 40 minutes. Scale bars represents 25 μm and 5 μm for inset.

Recent work from our lab demonstrated that CYRI-A and B are involved in macropinocytosis leading to the bulk internalisation of integrins (Le et al., 2021). Here, using B16-F1 *Cyri-b* KO cells, rescued with CYRI-B-p17-GFP and β1-integrin-mCherry we performed super-resolution live imaging and observed β1-integrin being internalised on vesicular structures surrounded by CYRI-B (Fig. 6d, Supp. Movie2) similar to what was previously reported in other cell types (Le et al., 2021).

We next asked if β1-integrin internalisation was affected in *Cyri-b* KO cells. Active β1-integrin antibodies were allowed to bind to the integrin extracellular domain and then to internalise for an allocated time before being removed from the extracellular surface. We observed a steady increase in the number of internalised vesicles containing β1-integrin in the control cells (Fig. 6e,f), which also resulted in a larger internal pools of vesicles containing β1-integrin (Fig. 6g). In contrast, the *Cyri-b* KO cells had significantly fewer and smaller β1-integrin containing vesicles internalised (Fig. 6e,f). Overall, we find a defect in β1-integrin internalisation in the *Cyri-b* KO B16-F1 cells resulting in an increase in active β1-integrin on the cell surface and in agreement with Le et al. (2021).

ERC1 is important for the internalisation of active integrins from the leading edge of migrating cells (Astro et al., 2014; Astro et al., 2016). Similar to the *Cyri-b* KO cells, the *Erc1* knockdown (KD) cells had more active β1-integrin present at the surface (Fig. 6h) and were much slower to internalise this into the cells (Fig. 6i,j). This confirms previous data that ERC1 promotes active integrin internalisation (Astro et al., 2014; Astro et al., 2016) and supports our hypothesis that depletion of ERC1 from the leading edge of *Cyri-b* KO cells contributes to the enlarged FA phenotype.

### *Cyri-b* loss prevents ERC1 localising near focal adhesion sites due to enhanced actin cytoskeletal tension

We further explored possible mechanisms by which CYRI-B depletion might enhance FAs and prevent ERC1 reaching the leading edge. As FAs form through the activation of integrins and mature under the influence of actin retrograde flow, we speculated that actin retrograde flow may be different in *Cyri-b* KO cells, disrupting normal adhesion maturation. As the *Cyri-b* KO cells form broad lamellipodia and have more active-Rac1 (Fort et al., 2018), we measured the actin retrograde flow in B16-F1 cells. Actin was marked in the lamellipodia tip by photoactivatable-GFP-Actin (PA-GFP-Actin) and over time we observed that there was no significant difference in the actin retrograde flow between control and *Cyri-b* KO cells (Fig. 7a,b, Supp. Movie 3). Therefore, we conclude that the enlarged FA in the *Cyri-b* KO cells are not likely caused by changes in actin retrograde flow.

**Figure 7:**
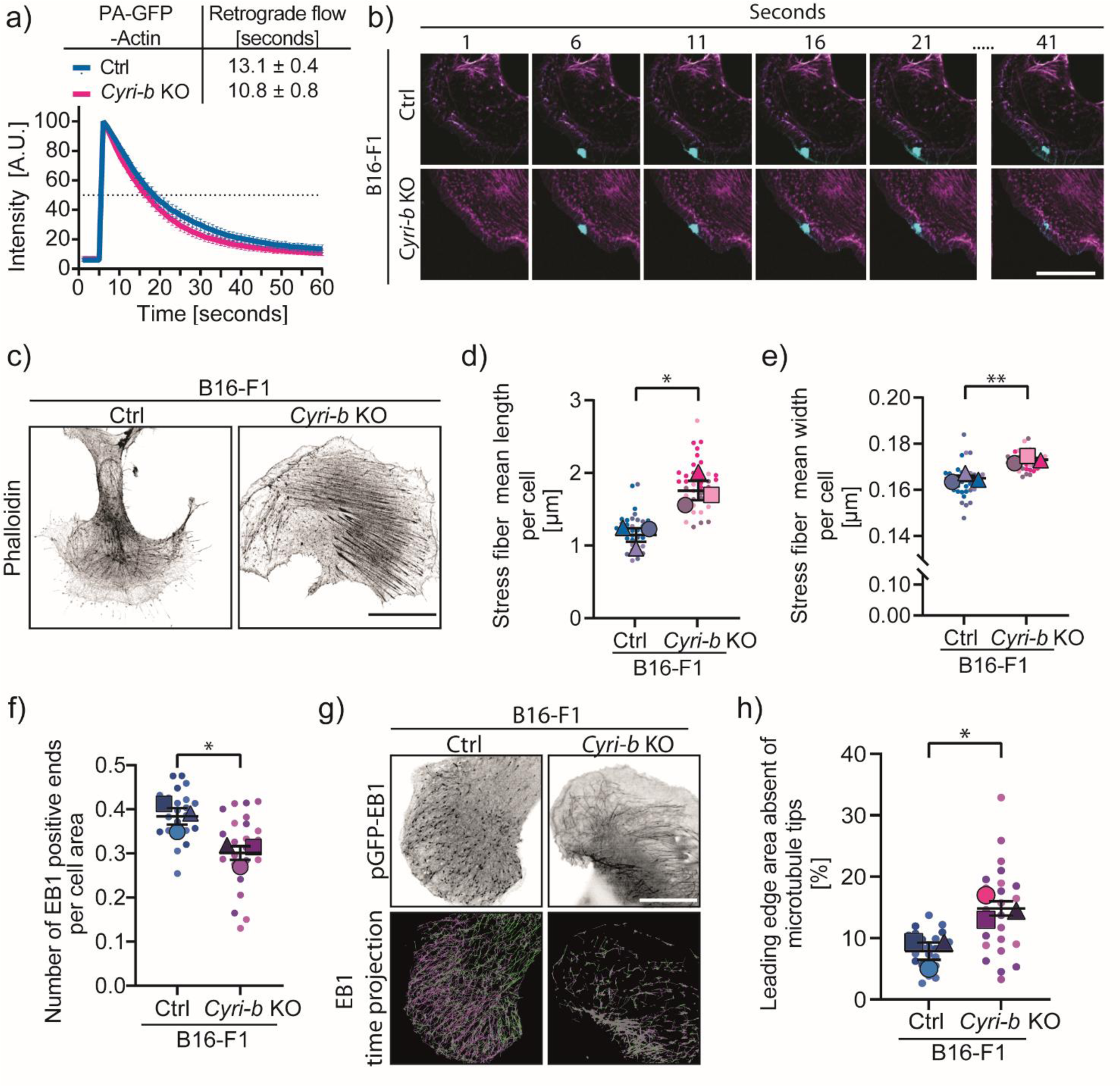
ERC1 trafficking is affected by reduced microtubule ends, which depend on normal actin dynamics and contractility. **a-b)** the actin retrograde flow was assessed in B16-F1 control and *Cyri-b* KO cells. **a)** The half-time from activating PA-GFP actin at the lamellipodia edge to flow into the lamella region of the cell. The peak in intensity correlates with photoactivation after 5 seconds. Intensity plot over time from 60 cells from 3 independent experiments where the error bars represent mean ± 95 % C.I. Average retrograde flow time is shown in the upper box ± S.D. **b)** Representative images of photoactivation of PA-GFP-Actin (cyan) and the actin cytoskeleton shown using LifeAct-TagRed (magenta) at various timepoints. Scale bar represents 20 μm. **c)** Representative images of stress fibers quantified using Phalloidin staining to highlight the F-actin cytoskeleton. Scale bar represents 25 μm. **d)** Average stress fiber length and **e)** average stress fiber thickness. 40 cells measured from 3 independent experiments. Error bars represent mean ± S.E.M., statistical significance determined using an unpaired two-tailed t-test. *P<0.05, **P<0.01. **f)** The number of EB1 positive microtubule ends normalised to cell area 25 cells measured from 3 independent experiments. Error bars represent mean ± S.E.M. in superplot format, statistical significance determined using an unpaired two-tailed t-test. *P<0.05. **g)** Representative images of *Cyri-b* KO B16-F1 cells expressing GFP-EB1 in greyscale (top) and a time projection (bottom) where magenta shows EB1 travel towards the leading edge and green as the EB1 travelling to the cytoplasmic region. Scale bar represents 25 μm. **h)** Quantification of the area at the leading edge without microtubules as a percentage of the cell area. 25 cells measured from 3 independent experiments. Error bars represent mean ± S.E.M. in superplot format, statistical significance determined using an unpaired two-tailed t-test. *P<0.05.

We noticed an increase in F-actin cables throughout the *Cyri-b* KO cells. This was not surprising, as mature FAs connect with actin stress fibers and regulate tension via Zyxin and α-actinin (Burridge and Guilluy, 2016). Quantitative image analysis revealed that the *Cyri-b* KO cells have longer and thicker actin stress fibers when compared to the control cells (Fig. 7c-e). Next, we asked whether the reduction of microtubule growth rates could be due to the contractile tension or steric hindrance from the strong actin stress fibers and/or a blockage from excessive actin accumulation at the leading edge of the cell. To answer this, we used either low dose treatment of Latrunculin A (LatA) to reduce actin assembly at the leading edge (Yarmola et al., 2000) or we treated the cells with low dose blebbistatin to inhibit myosin-II contractility (Martino et al., 2018). Both low dose LatA and blebbistatin treatment rescued the EB1 growth rates in the *Cyri-b* KO cells to that of control cells (Fig. 8a, Supp. Movie 4). Furthermore, these treatments also rescued FA sizes in the *Cyri-b* KO cells (Fig. 8b-d).

**Figure 8:**
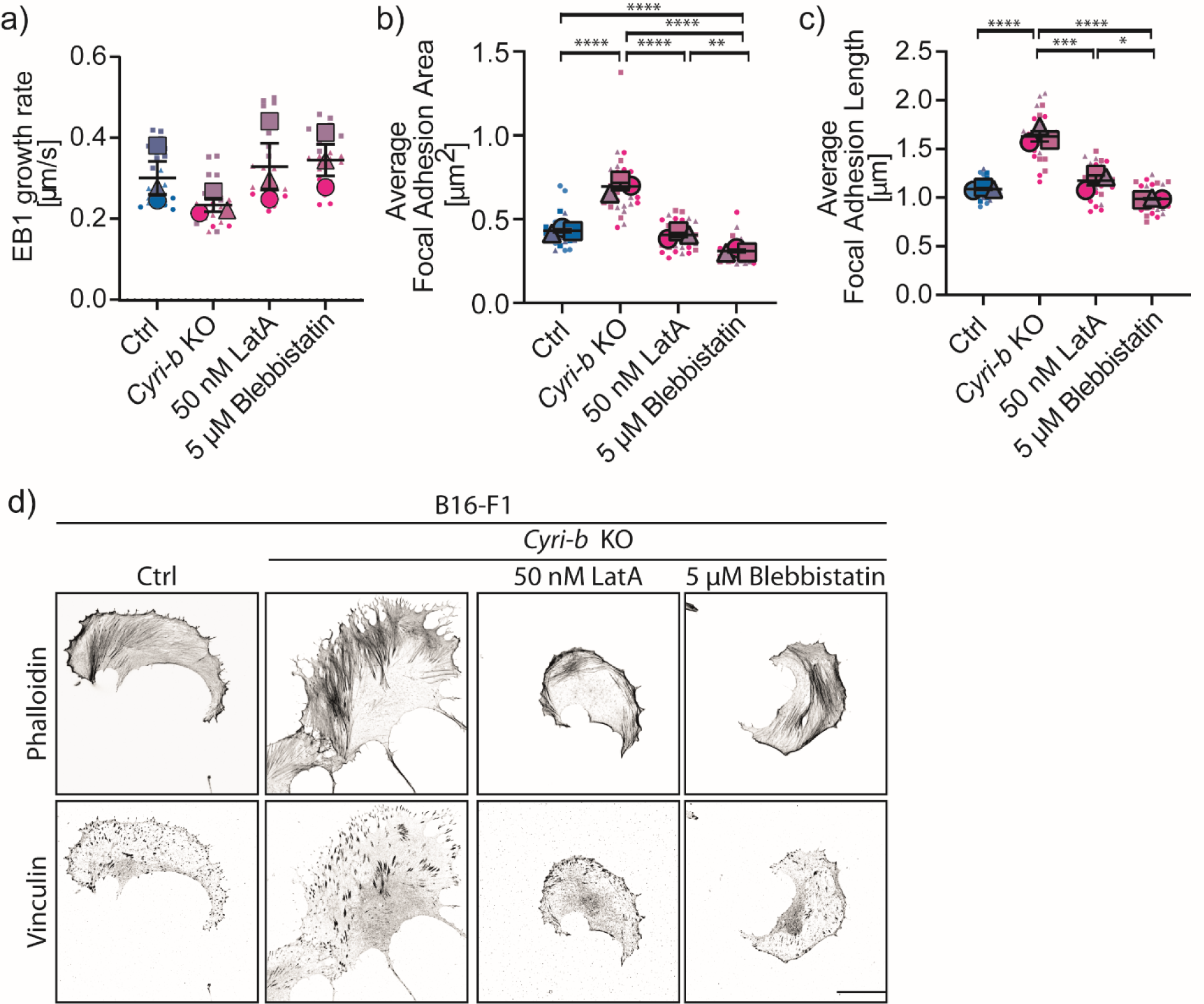
Loosening the actin tension and contractility restores normal microtubule growth rates and focal adhesion sizes. **a)** EB1 growth rates in B16-F1 control and *Cyri-b* KO cells with inhibitors. 25 cells were analysed over 3 independent experiments. Error bars represent mean ± S.E.M. in superplot format. Statistical significance measured by a 1-way ANOVA; No significance was not reached. **b-d)** Low dose chemical disruption to the actin cytoskeleton or cell contractility with 50nM LatrunculinA or 5 µM Blebbistatin, respectively. **b)** FA area and **c)** FA length in B16-F1 control or *Cyri-b* KO cells with inhibitors. 30 cells were analysed over 3 independent experiments. Error bars represent Mean ± S.E.M. in superplot format. Statistical significance measured by a 1 -way ANOVA, *P<0.05, **P<0.01, ***P<0.001, ****P<0.0001. **d)** Representative images of FA sizes in B16-F1 control and *Cyri-b* KO cells treated with 50 nM LatrunculinA or 5 µM Blebbistatin. Scale bar represents 25 μm.

Next, we looked at microtubule dynamics to see if microtubule positive end tracking was altered. The arrival of ERC1 is thought to displace the complex of FA proteins and allow the internalisation and recycling of integrins from the surface (Astro et al., 2016; Bouchet et al., 2016; Paradzik et al., 2020). Here, we used GFP-tagged EB1 (end-binding-1) to track the growth rates of microtubules. We observed a drastic reduction in the number of EB1 positive ends in the *Cyri-b* KO cells (Fig. 7f). Furthermore, by tracking EB1 movement at the tips, we determined that the microtubules in the *Cyri-b* KO cells did not reach the lamellipodia edge. This led to the *Cyri-b* KO cells having a larger area at their leading edges that was devoid of microtubules (Fig. 7g,h). Here, we conclude that a lack of microtubule plus ends tracking into the cell periphery could underly the reduced ERC1 localisation at the leading edge of cells and account for the reduced focal adhesion turnover we observed.

Overall, this suggests that the over-active actin cytoskeleton in the *Cyri-b* KO cells inhibits access of microtubule ends to the FA, preventing removal of β1-integrin by the ERC1/Liprin-α1/Kank complex. Taken together with our previous study showing how CYRI proteins function in integrin internalisation via macropinocytosis (Le et al., 2021), we conclude that actin dynamics and contractile function control access of microtubule ends to the leading edge of the cell. Microtubule access promotes the loosening up of FAs by ERC1/Liprin-α1, which allows integrin internalisation and normal recycling function (Fig. S3). Thus, the actin and microtubule cytoskeleton linkage are crucial for coupling of integrin trafficking with leading edge dynamics.

## Discussion

While CYRI proteins are known to regulate leading edge actin dynamics via Scar/WAVE complex and RAC1, very little is known about how they might crosstalk with nascent adhesions forming in lamellipodia. We previously found that depletion of CYRI proteins led to excess β1-integrin displayed on the cell surface, due to a reduction in internalisation via macropinocytic uptake (Le et al., 2021). However, it was unclear whether or how inhibition of integrin internalisation by macropinocytosis affected adhesion dynamics. Here, we find that depletion of CYRI-B enhances the size and changes the composition of focal adhesions, leading to enhanced maturation and a fibrillar elongated appearance. *Cyri-b* KO cells spread more rapidly than controls and show more rapid accumulation of proteins such as zyxin, that are hallmarks of mature adhesions (Zaidel-Bar et al., 2003). We initially speculated that adhesion turnover might be affected by the ability of CYRI to modulate RAC1 activation, but we found that RAC1 hyperactivation did not fully account for the phenotype of *Cyri-b* KO cells. We therefore set out to determine how CYRI-B regulates dynamic adhesion turnover.

To better understand the mechanisms for enhanced focal adhesion maturation in *Cyri-b* KO cells, we performed a Bio-ID screen to identify proteins in proximity to paxillin in focal adhesions of control vs knockout cells. Paxillin has one of the greatest numbers of protein binding partners within a FA and is ideal to use as the base for understanding changes in the adhesome (Chastney et al., 2020; Zaidel-Bar et al., 2007). We found multiple targets enriched in the focal adhesions of *Cyri-b* KO cells that suggested a role in mechanosensing, maturation and contractility. Hits included Shroom 2/4, which are implicated in contractility via RhoA activation (Simoes et al., 2014); pragmin, a pseudokinase that promotes RhoA activation via the small GTPase Rnd2 (Tanaka et al., 2006) tensin3, implicated in promoting oncogenesis and as a component of fibrillar adhesions (Atherton et al., 2021); vinexin and PAK2, both implicated in mechanotransduction and force production (Campbell et al., 2019; Kuroda et al., 2018) (Fig. S2c,d). We also found that ERC1, a protein implicated in internalisation of focal adhesion proteins (Astro et al., 2016) was enriched in the proximity of control adhesions over the knockouts.

Microtubule targeting to adhesions was originally shown to relax adhesions by Kaverina et al. (1999) and is thought to deliver proteins such as ERC1, which dock and displace adhesion proteins to allow internalisation. Due to its role in adhesion turnover, we followed up ERC1 and confirmed that it was indeed depleted from the leading edge of *Cyri-b* KO cells. Furthermore, depletion of ERC1 showed a similar phenotype to *Cyri-b* KO cells, supporting the idea that loss of CYRI-B impacts of focal adhesion turnover via ERC1. It remained an open question how loss of CYRI-B restricted ERC1 access to the cell leading edge. We reasoned that the excess actin assembly around the leading edge of *Cyri-b* KO cells might restrict access to the leading edge by the microtubule ends that were delivering ERC1. The enlarged adhesion sizes could also lead to positive feedback enhancing actin stress fibers and further obstructing ERC1 from accessing adhesion sites. We noticed a striking lack of EB1-positive microtubule ends tracking toward the periphery of many *Cyri-b* KO cells, supporting this hypothesis. Furthermore, if we lessened the actin network or the contractile myosin network with low doses of latrunculin-A or blebbistatin, we could rescue the delivery of microtubule ends to the periphery of the cell and rescue the effect of CYRI-B depletion.

While our data support the idea that CYRI-B loss promotes actin cytoskeletal changes that prevent microtubule- and ERC1-induced dynamic disassembly of focal adhesions, we acknowledge that our study has limitations. Firstly, we have not shown direct docking of ERC1 at focal adhesions, but rather leading-edge localisation that is disrupted in CYRI-B knockouts. Secondly, we did not detect a direct interaction between CYRI-B and ERC1, suggesting that the effect of CYRI-B deletion on ERC1 is indirect and likely due to cytoskeletal changes. We think that the most likely explanation for the effects of CYRI-B loss on focal adhesion dynamics is the combined effect of lack of targeting of microtubule tips to the leading edge of cells where nascent adhesions are forming with the previously described role of macropinocytosis of integrins (Le et al., 2021). Direct observation of ERC1 and integrin co-trafficking in normal and CYRI-B knockout cells would be needed to establish this mechanism, which awaits future studies.

Taken together, our results suggest that CYRI proteins enhance dynamic actin turnover at the leading edge of the cell to allow microtubule and ERC1 access to the leading edge to accelerate focal adhesion dynamics. Disruption of this turnover by depleting CYRI-B led to enhanced stability and maturation of focal adhesions, which feeds back positively to enhance stress fibers and recruitment of pro-contractility proteins to focal adhesions (Fig. S3). It will be interesting to know whether ERC1-mediated integrin internalisation is linked to macropinocytosis or whether these represent two separate and possibly additive mechanisms for mediating integrin internalisation from the cell surface.

## Materials and Methods

### Mammalian cell culture conditions

Mouse embryonic fibroblasts (MEFs) and mouse melanoma B16-F1 cells were maintained in Dulbecco’s Modified Eagles Medium (DMEM) supplemented with 10 % FBS, 2 mM L-glutamine at 37 °C, 5 % CO_2_. MEFs complete DMEM was supplemented with 1 mg ml^-1^ primocin. Cells were routinely tested for *Myocoplasma* contamination (MycoAlert; Lonza).

### Transfection of mammalian cell lines

*Cyri-b^fl/fl^*mouse embryonic fibroblasts were transiently transfected by electroporation (Amaxa, Kit T, program T-020) with 5 B16-F1 cells were plated on a 6-well plate and grown to 70 % confluency and later transfected with Lipofectamine 2000 following the manufacturer’s guidelines with 2-5 μg DNA.

### Genetic knockouts

Inducible knockout of *Cyri-b^fl/fl^* MEFs were generated by addition of 1 μM 4-hydroxytamoxifen (OHT) in the growth medium, with cells being split on day 2 and used in an assay on day 4 as described in Fort et al. (2018).

### Generation of Cyri-b KO B16-F1 cells

*Cyri-b* knockout in B16-F1 mouse melanoma cells were generated using the Cas9-GFP system and cell sorting. Specific gRNAs against mouse *Cyri-b* (ex3: CACCGGGTGCAGTCGTGCCACTAGT) were cloned into the sPs-U6-gRNA-Cas9 (BB)-2A-GFP vector (Addgene Plasmid #48138) (Ran et al., 2013). B16-F1 cells were transiently transfected with Cas9-GFP vectors and FACS sorted for GFP positive cells 36 hours after transfection. The empty sPs-U6-gRNA-Cas9 (BB)-2A-GFP vector was transiently transfected in B16-F WT cells as a control. Stable clones were isolated and tested for deletion of CYRI-B by Western blotting.

### siRNA knockdowns

*Erc1* was genetically knocked down in B16-F1 WT cells using specific siRNA oligonucleotides targeting Rab6ip (Erc1) (Qiagen; 1027416). The cells were transfected using Lullaby transfection reagent according to the manufacturer’s instructions with a pool of 10 nM of Mus musculus Rab6ip siRNA (2.5 nM each) or a matched concentration of control scramble siRNA (AllStars Negative siRNA, Qiagen; 1027281). The knockdown efficiency of ERC1 was determined by Western blotting using Mouse anti-ELKS antibody (Sigma; E4531).

### SDS-PAGE and western blotting

Cell lysates were collected on ice by scraping cells in RIPA buffer (150 mM NaCl, 10 mM Tris-HCl pH 7.5, 1 mM EDTA, 1 % Triton X-100, 0.1 % SDS, 1X protease and phosphatase inhibitors). The tubes were centrifuged for 10 minutes at 15,000 rpm and 4 °C. The lysate was transferred to a clean Eppendorf tube and protein concentration was measured using Precision Red.

40 μg of protein lysate was resolved on NuPAGE Novex 4-12 % Bis-Tris gels and transferred onto nitrocellulose membranes (Bio-Rad system). Membranes were blocked with 5 % BSA in TBS-T (10 mM Tris pH 8.0, 150 mM NaCl, 0.5 % Tween-20) for 20 minutes prior to overnight incubation with the primary antibody at 4 °C on a shaking incubator. Membranes were then washed three times for 5 minutes each in TBS-T. Membranes were incubated at room temperature for 1 hour with secondary DyLight conjugated antibodies 680 and 800 (ThermoFisher Scientific). The blots were washed again for 5 minutes in TBS-T three times before being imaged on the Li-Cor Odyssey CLx machine. Images were then analysed using the Image Studio Lite Version 5.2 and protein band intensities were calculated. These were then plotted in GraphPad Prism9 as a bar chart highlighting each repeat as a different shape and colour.

### Immunofluorescence analysis

Cells were plated onto sterile 13 mm glass coverslips that had been previously coated with either 10 μgml^-1^ Rat tail Collagen I (MEFs) or 10 μgml^-1^ laminin (B16-F1 cells). Cells were fixed with 4 % paraformaldehyde for 10 minutes at room temperature (RT). Coverslips were then washed three times with PBS before incubation with blocking buffer (0.05 % Triton X-100, 5 % BSA, PBS) for 15 minutes, with shaking. Primary and secondary antibodies were diluted in blocking buffer and incubated with the coverslips in a dark, humidified chamber for 1 hour. Coverslips were washed three times in PBS and once in MilliQ water before mounting with FluoromountG solution containing DAPI (Southern Biotech; 0100-01).

### Antibodies

Mouse anti-Vinculin (Sigma; clone hVIN-1), Rabbit anti-Vinculin (Sigma; 700062), Mouse anti-Zyxin (Abcam; ab50391), Rabbit anti-Zyxin (Sigma; HPA004835), Mouse anti-Talin1 8D4 (Sigma; T3287), Mouse anti-FAK (ThermoFisher Scientific; 34Q36), Rabbit anti-phospho-FAK (Y925) (CST; 3284S), Mouse anti-Paxillin (BD Bioscienses; 610052) and Rabbit anti-phospho-Paxillin (Y31) (ThermoFisher Scientific; 44-720G), Rabbit anti-β1-integrin (Cell Signalling Technologies; 4706), Rat anti-β1 subunit of VLA (Millpore; 1997), Rat anti-CD29 clone: 9EG7 (BD Pharmingen; 553715), Mouse anti-ELKS (Sigma; E4531), Rabbit anti-ERC1 (Atlas antibodies; HPA019523), Chicken anti-PPFIA1/Liprin α1 (Abcam; ab26192), Mouse anti-GFP (Abcam; Ab1218), AlexaFluor conjugated Phalloidins (ThermoFisher Scientific).

Western blot loading controls: Mouse α-Tubulin (Clone DM1A, Sigma; 9026) or Rabbit GAPDH (Cell Signalling Technologies; 14C10).

### Microscopy imaging

Fluorescent images were acquired using either; a Zeiss 880 confocal microscope with Airyscan using a Plan-Apochromat 63x/1.4 oil DIC objective lens and 405nm, 488nm, 561nm and 633nm laser lines. Raw images were acquired and Airyscan processing was performed using Zen Black version 2.3 SP1. Or a Zeiss 710 confocal microscope using an EC Plan-NEOFLUAR 40x/1.3NA Oil DIC and 405nm, 488nm, 561nm and 633nm laser lines running on Zen Black version 2011 SP7.

Images were processed using Fiji Version 1.53q.

### Focal adhesions

Cells were cultured as described above. The coverslips were fixed and stained with AlexaFluor_647_ Phalloidin and Mouse anti-Vinculin to measure cell area and FAs, respectively.

Z-stacked images were acquired using a Zeiss 880 confocal microscope with Airyscan using a Plan-Apochromat 63x/1.4 oil DIC objective lens and analysed using Fiji software. A maximum intensity projection (MIP) of the Z-stack image with 0.25 µm increments was performed, the FAs were enhanced using a Gaussian blur filter (2.0) and identified using ImageJ’s find maxima within tolerance. The output image from the ImageJ-derived maxima was overlaid onto a greyscale image of the FAs from the original file to indicate that the method can distinguish most FA proteins from the original image. Where erroneous structures were detected, manual deletion of the area was done before measurements. These were then measured using the Analyse Particles Plugin in Fiji to give FA area and length.

As an unbiased approach, we quantified morphological characteristics such as FA area using CellProfiler software (v2.4.0). Applying the CellProfiler pipeline as described in Cutiongco et al. (2020), where FAs were identified by vinculin staining. The individual adhesions were measured for their area and displayed as a frequency graph using Orange 3.30.2 software.

### Focal adhesion ratios

B16-F1 cells were grown on coverslips as described. The coverslips were fixed and stained with either Mouse anti-Vinculin or Rabbit anti-Vinculin antibodies as a standard to normalize all other FA antibodies against, such as Rabbit anti-Zyxin, Mouse anti-Talin1, Mouse anti-FAK, Mouse Paxillin and Rabbit anti-phospho-Paxillin (Y31). Images were acquired as above, and the Fiji Plot Profile tool was used to measure the fluorescence intensity over the FA from the lamellipodia tip going into the cytosol. The fluorescence intensity was first normalized where the highest intensity reading for each antibody was given the 100 % value and the subsequent values a percentage of the highest. As all the FAs were of varying lengths, dividing the intensity reading into 100 equal parts normalized the plot profile. These were then plotted using GraphPad Prism to generate a heatmap. The graphical output provides an indication of the complexity of the FAs and where each protein is presented as the abundance from the periphery (tip) to cytosol (rear) of the FA. More than 50 FAs were imaged for each antibody pairing.

### β1-Integrin area

B16-F1 cells were plated on laminin coated coverslips and left to spread. The coverslips were fixed and stained for Rat anti-β1 integrin and AlexFluor568 phalloidin. Z-stacked images with 0.25 µm increments were captured using a Zeiss 880 confocal microscope with Airyscan using a Plan-Apochromat 63x/1.4 oil DIC objective lens. In Fiji, a Gaussian filter was applied to the max projected images to reduce background and highlight the integrin signal. As there was a saturated signal in the cytoplasmic region around the nucleus that would affect the quantifications, we removed this region and focused the analysis on the lamella and lamellipodia regions of the cell. These were then measured using the Analyse Particles Plugin in Fiji to give β1 integrin area.

### Image-based Integrin internalisation assay

This assay aims to quantify the internalisation of β1 integrin over time. B16-F1 cells were grown on laminin coated coverslips overnight as described above. The next day, cells were washed once with ice-cold PBS and incubated with Rat anti-β1-integrin antibody clone 9EG7 diluted in ice cold Hank’s Balanced Salt Solution (HBSS) for 1 hour on ice in a dark humid chamber.

Integrin internalisation was induced by the addition of 1 ml of pre-warmed DMEM complete and quickly transferred to a 37 °C incubator for specified times (10, 20, 40 minutes). After the allotted time, the coverslips were washed once with ice-cold PBS and incubated for 5 minutes in stripping buffer (0.2 M acetic acid, 0.5 M NaCl, pH 2.5) to remove all extracellular bound antibody. The coverslips were washed a further time in ice-cold PBS and fixed with 4 % PFA.

For the controls, a total β1-integrin integrin measurement was taken, whereby the cells were fixed prior to any antibody treatment. A second control to determine the efficiency of antibody stripping after incubation was the 0-minute coverslip. Here, after incubation with the β1 integrin antibody, the coverslips were kept on ice, washed with the stripping buffer and not allowed to internalise. This control should not have any internalised β1-integrin.

After fixation, the coverslips were subjected to the immunofluorescence protocol as described above with only the blocking and permeablising step before the addition of the secondary antibody against Rat.

For the image acquisition, a Z-stack image was taken with a Zeiss 880 with AiryScan module using the Plan-Apochromat 63x/1.4 oil DIC objective lens. In Fiji, a maximum projection image was generated from Z-stacked image with 0.16 µm increments, a Gaussian blur of 2.0 was applied to the image to reduce background noise. Manual thresholding was applied to the images and using the Analyse Particle plugin of Fiji to quantify the number of internalised β1-integrin dots and the area of those dots normalised to the cell area. 40 fields of view were analysed from each condition over 4 independent experiments.

### CYRI-B GFP positive vesicles containing β1-integrin

B16-F1 cells were transiently transfected with CYRI-B-p17-GFP and mCherry-β1 integrin (Addgene plasmid #55064) and plated on laminin coated glass bottom dishes. Images were acquired using a Zeiss 880 confocal microscope with Airyscan using a Plan-Apochromat 63x/1.4 oil DIC objective lens with a 37 °C heated incubator, perfused with 5 % CO_2_. Images were acquired every 10 seconds for

5 minutes. Images were processed using Fiji software and a 2.5 µm line through the vesicles was drawn and a plot profile intensity was captured. The intensities were then normalized where the brightest intensity was given a 100 % value with the other values as a percentage of the highest value. Each vesicle was then averaged and displayed on a line graph using GraphPad Prism.

### Focal adhesion formation and maturation

B16-F1 cells were trypsinised for 2 minutes and resuspended with DMEM complete and adjusted to 1x10^5^ cells per ml, with 500 µl added to each coverslip coated with laminin before being placed in the incubator for the specific times (10, 30 mins, 1 and 3 hours). The coverslips were gently fixed with 4 % PFA to preserve the cells that had weakly attached. The coverslips were stained with mouse anti-Paxillin as an early adhesion marker and Rabbit anti-Zyxin as a marker for more mature FAs and AlexaFluor_647_ Phalloidin for cell area.

Z-stack images were acquired using a Zeiss880 microscope with AiryScan module, Plan-Apochromat 63x/1.4 oil DIC objective lens 405nm, 488nm, 561nm and 633nm laser lines. The max intensity projection images from 9 slices at 0.2 µm increments were analysed using Fiji and both Paxillin and Zyxin area and length was quantified over time to distinguish adhesion formation from nascent to mature FAs as described above. Data are presented from 3 independent experiments in superplot format.

### Focal adhesions turnover

B16-F1 cells were transiently transfected with pEGFP-Paxillin (Addgene plasmid #15233) as described above and plated onto 35 mm glass-bottom Ibidi dishes coated with laminin. Short movies of 1 frame per minute for 30 minutes were obtained using the 488 nm laser on the Zeiss LSM 880 confocal microscope with Airyscan module using a Plan-Apochromat 63x/1.4 oil DIC objective lens at 37 °C and 5 % CO_2_. Raw images were acquired and Airyscan processing was performed using Zen Black version 2.3 SP1. Time-lapse movies were processed using Fiji software 1.53q, where the image sequences were stabilized using the Fiji plugin Image stabilizer and a Gaussian blur 2.0 was applied to the image to highlight the focal adhesions. If there were more than one cell imaged in a field of view, then this was edited to focus only on one cell throughout the duration of the movie. The movies were submitted to the Focal adhesion analysis server (http://faas.bme.unc.edu/) (Berginski and Gomez, 2013) where a threshold of 2.5 units was maintained across all image sets and positive structures or 15 pixels^2^ that last for at least 5 consecutive frames were quantified as being a focal adhesion. Assembly and disassembly rates are presented as rates from the FAAS. Data presented from 3 independent experiments in superplot format.

### xCELLigence cell spreading

E-plate 16 were coated with laminin overnight and equilibrated with DMEM complete for 30 minutes prior to imaging at 37 °C. Cells were harvested and adjusted to 5x10^3^ per well. The cells were seeded in technical quadruplicate and the plate was immediately transferred to the Acea RTCA DP xCELLigence machine maintained at 37 °C, 5 % CO_2_. Cell index was measured at 5-minute time intervals for 8 hours and readings were averaged for each condition. The impedance between the electrodes and cells determined cell index over time. Quadruplicate readings were taken for each condition. Data are presented as the average impedance from 3 independent replicates as described in Whitelaw et al. (2020).

### BioID-Paxillin

B16-F1 cells were stably transfected with GFP-BirA*-Paxillin (kindly gifted by Dr. Ed Manser, Institute of Molecular and Cell Biology, Singapore) and a pPuro empty vector. The cells were first selected with puromycin (2 µg/ml) and then after cell survival, the cells were then FACS sorted for low to mid-range GFP expression. Cells were plated on 15 cm laminin coated dishes and left to grow to around 50 % confluence overnight. The following day, the dishes were treated with either 50 µM biotin ligase or DMSO for another 16 hours.

For purification of the biotinylated proteins, the dishes were washed twice with ice cold PBS, with the cells being scraped off the dish in 300 µl lysis buffer (50 mM Tris pH 7.2, 1 % NP-40, 0.1 % SDS, 500 nM NaCl, 10 mM MgCl_2_, 5 mM EGTA, pH 7.5) and incubated in the tube for 10 minutes prior to centrifugation (20 minutes, 15,000 rpm, 4 °C). The protein was then transferred to a clean tube and quantified using PrecisionRed (Cytoskeleton; ADV02-A) at OD_600_.

For each condition, 1.5 mg of protein was made to a volume of 500 µl in lysis buffer and added to 500 µl Tris-Cl pH 7.4 for a total 1 ml volume. This was then added to 50 µl Pierce NeutrAvidin Agarose bead slurry (ThermoScientific; 29200) that was pre-washed twice with 250 µl lysis buffer. The tubes were then incubated overnight at 4 °C on a rotating block. The next day, the tubes were spun at 1500 rpm, 4 °C for 1 minute and resuspended in Wash buffer 1 (2 % SDS). The tubes were then rotated for 8 minutes at room temperature due to high SDS content in Wash buffer 1. The Wash buffer 1 step was repeated and after the spin, the beads were resuspended in 1 ml Wash buffer 2 (0.1 % Sodium deoxycholate, 1 % NP-40, 1 mM EDTA, 500 mM NaCl, 50 mM HEPES, pH 7.5). The mixture was rotated for 2 minutes, then spun at 1500 rpm and resuspended with 1 ml Wash buffer 3 (0.5 % sodium deoxycholate, 0.5 % NP-40, 1 mM EDTA, 250 mM LiCl, 10 mM Tris-Cl, pH 7.4). The tubes were rotated for a further 2 minutes and after the spin, resuspended with 1 ml Tris-Cl. This wash step was repeated with 1 ml Tris-Cl and the beads were spun down. As much of the liquid was removed as possible, for mass-spectrometry analysis.

For initial proof of concept, 2X sample buffer was added to the beads after the wash steps and heated to 100 °C for 10 minutes. This was then run for western blot analysis and blots were probed using anti-streptavidin-HPR (ThermoScientific; N100).

### Sample preparation

Agarose beads were resuspended in a 2M Urea and 100mM ammonium bicarbonate buffer and stored at -20°C. On-bead digestion was performed from the supernatants. biological replicates (n=7) were digested with Lys-C (Alpha Laboratories) and trypsin (Promega) on beads as previously described (Hubner et al., 2010).

### MS Analysis

Peptides resulting from all trypsin digestions were separated by nanoscale C18 reverse-phase liquid chromatography using an EASY-nLC II 1200 (Thermo Scientific) coupled to an Orbitrap Fusion Lumos mass spectrometer (ThermoScientific). Elution was carried out at a flow rate of 300 nl/min using a binary gradient, into a 50 cm fused silica emitter (New Objective) packed in-house with ReproSil-Pur C18-AQ, 1.9 μm resin (Dr Maisch GmbH), for a total run-time duration of 135 minutes. Packed emitter was kept at 50 °C by means of a column oven (Sonation) integrated into the nanoelectrospray ion source (ThermoScientific). Eluting peptides were electrosprayed into the mass spectrometer using a nanoelectrospray ion source. An Active Background Ion Reduction Device (ESI Source Solutions) was used to decrease air contaminants signal level. The Xcalibur software (Thermo Scientific) was used for data acquisition. A full scan over mass range of 350–1550 m/z was acquired at 60,000 resolution at 200 m/z. Higher energy collisional dissociation fragmentation was performed on the 15 most intense ions, and peptide fragments generated were analysed in the Orbitrap at 15,000 resolution.

### MS Data Analysis

The MS Raw data were processed with MaxQuant software (Cox and Mann, 2008) version 1.6.3.3 and searched with Andromeda search engine (Cox et al., 2011), querying SwissProt (UniProt, 2019) Mus musculus (62094 entries). First and main searches were performed with precursor mass tolerances of 20 ppm and 4.5 ppm, respectively, and MS/MS tolerance of 20 ppm. The minimum peptide length was set to six amino acids and specificity for trypsin cleavage was required. Cysteine carbamidomethylation was set as fixed modification, whereas Methionine oxidation, Phosphorylation on Serine-Threonine-Tyrosine, and N-terminal acetylation were specified as variable modifications. The peptide, protein, and site false discovery rate (FDR) was set to 1 %. All MaxQuant outputs were analysed with Perseus software version 1.6.2.3 (Tyanova et al., 2016).

Protein abundance was measured using label-free quantification (LFQ) intensities reported in the ProteinGroups.txt file. Only proteins quantified in all replicates in at least one group, were measured according to the LFQ algorithm available in MaxQuant (Cox et al., 2014). Missing values were imputed separately for each column, and significantly enriched proteins were selected using a permutation-based t-test with FDR set at 5% or a cut-off at p-value 0.05.

Network of DTXs proteins interactors was generated from LFQ intensities using the Hawaii plot functionality in Perseus (Rudolph and Cox, 2019). Network of DTXs proteins interactors was generated from LFQ intensities using the Hawaii plot functionality in Perseus (Shannon et al., 2003) for network visualisation

### GFP-Trap

Transiently transfected B16-F1 cells expressing GFP or CYRI-B-p17-GFP were washed twice with PBS on ice and scraped with 400 μl of lysis buffer [25mM Tris HCl, pH7.5, 100mM NaCl, 5mM MgCl_2_, 0.5% NP-40, Protease and phosphatase inhibitors]. Lysates were kept on ice 30 minutes and thoroughly mixed every 10 minutes. Soluble proteins were collected after a 10 minute centrifugation step at 15000 rpm and protein concentration was measured using PrecisionRed (Cytoskeleton; ADV02). 1.5 mg of protein was mixed with 25 μl of pre-equilibrated GFP-Trap_A beads (ChromoTek) and incubated for 2 hours at 4°C with gentle agitation. Beads were then washed 3 times with 500 μl of wash buffer [100mM NaCl, 25mM Tris-HCl pH7.5, 5mM MgCl2].

To test for ERC1 interaction, 2X sample buffer and 2X reducing agent was added to the beads after the wash steps and heated to 100 °C for 10 minutes. This was then run for western blot analysis and blots were probed using anti-ERC1 (Sigma).

### ERC1 and Liprin localisation

B16-F1 cells were plated onto coverslips as above, fixed and stained with either Rabbit anti-ERC1 or Chicken anti-Liprin α1 and Alexa Fluor_647_ Phalloidin. Images were acquired using a Zeiss 710 confocal microscope and EC Plan-NEOFLUAR 40x/1.3NA Oil DIC objective lens. The images were processed using Fiji software and the cells were scored for either a membrane or a more diffuse localization and presented as a percentage. Membrane localization was deemed positive when there was a tight localisation around the leading edge of the cell. Diffuse signals had no distinct localization anywhere in the cell and presented similar to a non-specific staining. For the line graph, a 3 μm line and subsequent plot profile of fluorescence intensity from the cell edge into the cytosol was taken. The fluorescence signals were averaged and plotted to represent both control and *Cyri-b* KO cells with either a membrane or diffuse localization.

### Actin photoactivation - Retrograde flow

Photoactivation of actin and retrograde flow analysis was conducted as described in Papalazarou et al. (2020). Briefly, B16-F1 cells were transiently transfected with LifeAct-TagRed and PA-GFP-Actin (Addgene #57121) as described above. Imaging was conducted on a Zeiss 880 confocal microscope using a Plan-Apochromat 63x/1.4 oil DIC objective lens. The PA-GFP-Actin and LifeAct-TagRed were monitored with 488 nm and 568 nm lasers respectively. A single pulse with a 405 nm laser (pulse length t=0.5 seconds) obtained photoactivation of actin at the ROI. Acquisitions were taken every second for 60 frames with an initial 5 seconds to obtain baseline GFP intensity prior to activation. Data presented as the means from 3 independent experiments in a time decay graph.

### Stress fiber quantification

The B16-F1 cells were plated onto coverslips coated with laminin and incubated overnight at 37 °C and 5 % CO_2_. The coverslips were fixed and stained with AlexaFluor_647_ Phalloidin as described above. Z-stacked images obtained from a Zeiss880 microscope with AiryScan module, Plan-Apochromat 63x/1.4 oil DIC objective lens and 405nm and 633nm laser lines for DAPI and Phalloidin, respectively. Images were processed using the macro to max project the z-stack, highlight the stress fibers with a Difference of Gaussians threshold and Ridge Detection to identify and quantify stress fibers as described in Whitelaw et al. (2020). Data presented from 3 independent experiments.

### Microtubule ends

pGFP-EB1 (Addgene plasmid #17234) was transiently transfected into the B16-F1 control and *Cyri-b* KO cells and imaged live on a Zeiss 880 microscope with Airyscan with a Plan-Apochromat 63x/1.4 oil DIC objective lens with the 488nm laser at 1 image per second for 120 seconds. Image analysis was conducted using Fiji software to threshold for the EB1 microtubule tips. This number was then divided by the cell area.

Tracking of the EB1 positive tips was done using Fiji plugin TrackMaxima (IJ2). With setting the threshold to 8.0 and blur to 4.0. EB1 was tracked throughout the movie where the EB1 was in focus for at least 10 frames.

To measure the area of the lamellipodia absent of microtubules, the above movies were time projected using the Fiji TrackMaxima (IJ2) software. The whole cell area in the field of view was thresholded and used as a mask. The time projected EB1 tracks were used as a mask for how far the microtubules have travelled to the leading edge. The EB1 track mask was subtracted from the whole cell area mask to obtain an area devoid of microtubules at the leading edge of the cell. This devoid area was normalised as a percentage of the total area of the cell.

### Chemical inhibitors

Low dose LatrunulinA (Merck; L5163) and blebbistatin (Sigma; B0560) were used to disrupt the actin cytoskeleton and reduce cell contractility, respectively. Serial dilutions of the drugs or DMSO were added to B16-F1 *Cyri-b* KO cells to determine the concentration at which the cells were still able to form lamellipodia and show healthy morphological features. We established that treatment with either 50 nM LatA or 5 µM blebbistatin for 20 minutes prior to imaging was sufficient to rescue the phenotypes of the *Cyri-b* KO cells.

### Statistics and reproducibility

All datasets were analysed using GraphPad Prism version 9.3.1. Datasets were tested for normality and then analysed using the appropriate statistical test, as described in each figure legend. Where appropriate, SuperPlots were used (Lord et al., 2020). For this, each individual value was colour coded according to the experiment and the mean of each experiment were overlaid with larger symbols, also colour coded to experimental day. The statistical analysis was done on the experimental means and presented with SEM. Significance levels rejecting the null hypothesis are represented above figures where: NS P>0.05, * P<0.05 *, **P<0.01, *** P<0.001 and **** P<0.0001. Where significance was not reached, nothing was added above the graphs.

## Acknowledgments

We would like to thank Dr Ed Manser (Institute of Molecular and Cell Biology, A*STAR) and Dr Susan Farrington for sharing the GFP-BirA*-Paxillin BioID and turboGFP-Shroom2 construct with us, respectively. We thank the Machesky and Insall lab members for technical advice and discussions. We thank the Beatson Advanced Imaging Resource (BAIR), Margaret O’Prey, John Halpin and Tom Gilbey for their help with confocal microscopy and image analysis and flow cytometry, respectively. We thank the Beatson central service and molecular services.

